# Global RNA sequencing reveals enhanced photosynthetic activity and auxin response in *Brassica napus* treated with *Pseudomonas chlororaphis* PA23

**DOI:** 10.1101/2020.07.30.228635

**Authors:** Joey C Wan, Michael G Becker, Ayooluwa Bolaji, Philip L Walker, Emma Gray, Teresa R de Kievit, WG Dilantha Fernando, Mark F Belmonte

## Abstract

Plant growth promoting bacteria (PGPB) are a growing subset of agricultural adjuncts which can be used to increase crop yield and plant productivity. Although substantial research has been conducted on the metabolites and active molecules secreted by PGPBs, relatively little is known about their effects on the global transcriptome of the host plant. The present study was carried out to investigate changes in the gene expression landscape of early vegetative *Brassica napus* following treatment with *Pseudomonas chlororaphis* PA23. This PGPB was isolated from the soybean rhizosphere and has been extensively studied as a biocontrol agent. However, little is known about its effects on plant growth and development. Using a combination of RNA-sequencing and physiological analyses, we identified increased abundance of mRNA transcripts associated with photosynthesis and phytohormone response.

Phenotypically we observed increased photosynthetic rates and larger root and shoot systems in *B. napus* following *P. chlororaphis* PA23 treatment. Lastly, we identified auxin production by *P. chlororaphis* PA23 which likely contributes to changes in gene expression and the observed phenotypic differences in root and shoot structures. Together, the results of our study suggest that PA23 is a potent plant growth promoting agent with the potential for field applications as an agricultural adjunct.

## Introduction

Plant associated soil bacteria are a group of microbes that reside within or on the surface of plant roots and can have profound effects on plant health. These microbes can further be divided into two groups: plant growth promoting bacteria (PGPB) which impact growth through direct interactions with the plant, or biocontrol bacteria (BCB) which indirectly enhance growth by inhibiting harmful pathogens [1,2]. PGPBs have been shown to affect gene expression, phytohormone signalling, and mobilize soil nutrients into the plant [3]. The Pseudomonads are a group of bacteria that are universally found in agricultural settings.

Members of this genus have been studied extensively as both biocontrol and plant growth promoting agents [4–6]. For example, *Pseudomonas fluorescens* B16, promotes plant growth through the synthesis and secretion of the active molecule pyrroloquinoline which increased tomato plant height, flower number, and total fruit weight [7]. Additionally, treatment of wheat with a combination of *Pseudomonas* -FAP2 and *Bacillus licheniformis* resulted in increased vegetative growth, chlorophyll content, transpiration rates, and net photosynthetic rates [8].

Changes in vegetative growth, chlorophyll, and photosynthesis has been associated with the production and secretion of bacterial metabolites that are analogous or similar to phytohormones [9]. In addition, PGPBs are also capable of synthesizing plant growth promoting hormones; for example, bacterial auxin has been extensively studied as a plant growth promoting metabolite which modulates plant phytohormone signalling and alters root and shoot development [10]. Although the mechanisms of auxin induced growth are well understood, further work is required to determine the mode of action for potential PGPB due to differences in auxin synthesis rates and the effects of varying auxin concentrations on plants [11].

Other members of the pseudomonads, including strains of *P. fluorescens* and *P. chlororaphis*, can function as biocontrol agents to directly inhibit the growth of fungal, bacterial, and eukaryotic pathogens through the action of secreted compounds [12–16]. *P. chlororaphis* strain PA23, a microbe originally isolated from the soybean rhizosphere, is a potent biocontrol agent against fungal pathogens and parasitic nematodes [17,18]. Previous work has identified the mode of action for PA23 mediated biocontrol in the *Brassica napus-Sclerotinia sclerotiorum* pathosystem; specifically, PA23 directly inhibited pathogen growth through the activity of pyrrolnitrin, phenazine, and hydrogen cyanide [19,20]. Furthermore, PA23 primed plant pathogen defense systems resulting in enhanced tolerance to fungal infection [15]. However, little is known about *P. chlororaphis* plant growth promotion and few studies have investigated the interaction between *P. chlororaphis* and plant species, such as *B. napus*, that do not harbour a robust root microbiome. Given that the *P. chlororaphis* PA23 genome is sequenced, the secondary and secreted metabolites involved in this interaction can be predicted. Functional compounds include siderophores, hydrogen cyanide, and auxin, produced through the indole acetamide pathway, which may act to enhance plant growth while suppressing harmful pathogens [17,21].

Despite recent advances in sequencing technologies, we have yet to fully understand the plant growth promoting effects of pseudomonads and in particular the effects of *P. chlororaphis* PA23 on early vegetative growth in *B. napus*. Given that plant growth is governed by multiple factors including gene activity, a holistic understanding of the gene expression landscape is required to fully understand the putative mechanism of *P. chlororaphis* PA23 plant growth promotion. In our study, we used a combination of RNA-sequencing and physiological experiments to investigate how *P. chlororaphis* PA23 mediates changes in plant growth at the genetic and physiological level. Specifically, we identified elevated expression of genes associated with photosynthesis, nutrient uptake, and phytohormone signalling, which may contribute to increased vegetative growth rates of *B. napus* seedlings. Together, the results of our study provide a deeper understanding of the genetic mechanisms that govern *P. chlororaphis* plant growth promotion and contribute to the growing body of evidence for the use of PGPBs as an agricultural adjunct.

## Methods

### Plant and bacterial materials

*B. napus* cv. Westar seedlings were grown in growth chambers with a 16-hour photo period (21^°^C light period, 16^°^C dark period, and 150 µmol/m^2^/s). Ten plants per treatment per time point were grown in Sunshine Mix (SunGro Horticulture, Agawam, MA) in plastic pots (8 cm x 12 cm x 6 cm). Each pot was treated with 25 mL of *P. chlororaphis* PA23 at a concentration of 1⨯10^9^ cfu/mL or 25 mL of lysogeny broth (Difco Laboratories, Detroit, MI). Bacterial cultures were prepared from a frozen stock of PA23 stored in 10% skim milk solution (ThermoFisher, Waltham, MA). PA23 was grown on lysogeny broth agar plates (Difco Laboratories, Detroit, MI) at 28^°^C for 24 hours. Isolated colonies were used to inoculate lysogeny broth and incubated for 24 hours in a rotary shaker at 28^°^C and 250 rotations per minute. Cell pellets were formed through centrifugation at 5000 rpm for 10 minutes. Pellets were washed with sterile 0.9% saline, resuspended in lysogeny broth, and adjusted to a final concentration of 1⨯10^9^ cfu/mL (OD_600_ = 1) [15].

*Arabidopsis thaliana Col-0* seeds were surface sterilized with ethanol and germinated on half-strength Murashige and Skoog agar media (2.2 g/L MS vitamins, 1% sucrose, 0.8% agar, pH 5.8). Agar plates were incubated vertically under controlled conditions with an 8-hour photo period (21^°^C light period, 16^°^C dark period, 150 µE/m^2^/s). PA23 suspensions were prepared as previously described. Seven days after germination, Arabidopsis seedlings were treated with 2000 cfu of PA23 or mock-treated with 0.9% saline. Root architecture was visualized seven-days after bacterial exposure with the Leica M80 Dissecting microscope and images were captured with the Leica Application Suite.

We measured IAA production by *P. chlororaphis* PA23 using colorimetric spectrophotometry as described in Gordon and Weber [22] and Gang et al [23]. *P. chlororaphis* cultures were grown in 5 mL of nutrient broth in an incubating shaker at 28^°^C and 180 rpm. This culture was used to inoculate nutrient broth amended with L-tryptophan (1 g/L); which was incubated in the dark at 28^°^C and 180 rpm for 44 hours. After which, a 1.5 mL aliquot of the culture was centrifuged for 5 minutes at 16300 g to pellet cellular debris. The supernatant was removed and mixed with 1 mL of Salkowski’s reagent [24]. The reaction was carried out for 30 minutes in the dark at 30^°^C. Absorbance was measured at 546 nm and concentrations were determined with a linear regression against a standard curve containing indole-3-acetic acid.

### cDNA library synthesis and computational analyses

Total RNA was collected from two biological replicates with each replicate consisting of at least 5-10 *B. napus* plants treated with PA23 or lysogeny broth at 24 hours post treatment (hpt) and 7 days post treatment (dpt). RNA was isolated with PureLink Plant RNA Reagent (Ambion, ThermoFisher, Waltham, MA) and DNA contamination was removed with the TURBO DNA *free*^™^ (Ambion, ThermoFisher, Waltham, MA) kit according to the manufacturer’s recommendation. mRNA quality was determined with an RNA nanochip on the Agilent 2100 Bioanalyzer system (Agilent, Santa Clara, CA). cDNA libraries were prepared at Génome Quebec (Montreal, QC, Canada) with the Ultra II NEB DNA Library Prep Kit for Illumina (New England Biolabs, Ipswich, MA). Library quality and size distribution was determined on a high-sensitivity DNA chip on the Agilent 2100 bioanalyzer system. One hundred base pair single-end RNA sequencing was carried out on the Illumina HiSeq 2000 platform with a multiplex value of 16. Raw sequencing reads were trimmed, quality checked, and aligned according to the methods described in Becker et al 2017 [25]. Data clustering was performed on averaged raw counts with the PVClust package of R-Studio. Differentially expressed genes were identified with CuffDiff and the output was used with Venny, an online venn-diagram tool (https://bioinfogp.cnb.csic.es/tools/venny/), to identify treatment specific and co-expressed genes. Specific gene lists from Venny were used for GO term enrichment with SeqEnrich and terms were considered statistically enriched at P < 0.05 [26].

All sequencing data for this study are available online through NCBI accession GSE154315.

### Targeted real-time quantitative PCR

RNA was isolated from the same *B. napus* tissues as described above. Reverse transcription was carried out with the Maxima First Strand cDNA synthesis kit (ThermoFisher, Waltham, MA) according to the manufacturer’s protocol. Directed quantitative PCR was carried out on the Bio-Rad CFX Real-Time System with SYBR Green SuperMix (BioRad, Hercules, CA) as per the manufacturer’s instructions in a total reaction volume of 10 µL. Conditions for the reactions were as follows: 95^°^C for 3 minutes, 44 cycles of 95^°^C for 30 seconds, 58^°^C for 30 seconds, and 72^°^C for 30 seconds. Melt curves, 0.5^°^C increments over a range of 55^°^C to 95^°^C, for each gene target was performed to identify splice variants, primer dimers, and off-target amplification. A list of primer sequences used in these experiments is given in Table A in S1 text. Relative transcript abundance was determined using the ΔΔC_t_ method, normalizing to the endogenous housekeeping gene β-*Actin* and using lysogeny broth treated *B. napus* as the reference sample [27]. Results presented in Figure A in S1 text are based on two experimental repeats of two biological replicates with each replicate consisting of at least five *B. napus* tissue systems (Data in S3 Table). One-way ANOVA tests were performed with each gene to determine significant fold changes between treatment conditions (P < 0.05).

### Photosynthetic pigment extraction and quantification of photosynthetic rates

*B. napus* leaf tissue was flash frozen and homogenized in liquid nitrogen. At least 10 mg of 14-day old canola leaf tissue was placed into vials containing 10 mL of 80% acetone (20% v/v 0.2M Tris-HCl pH 8.0). Vials were stored at -20^°^C for 96 hours or until the pigments were cleared from the leaves. Pigment concentrations were determined with the Beer-Lambert Law and absorbance values at 663 nm and 645 nm [28]. Student’s t-tests were performed to identify significant differences in chlorophyll a and chlorophyll b between the mock-treated and PA23-treated conditions. Infrared gas analysis (IRGA) equipment was used to determine relative photosynthetic rates. Ambient CO_2_ levels were measured prior to introduction of photosynthetic tissues and used as the zero value. *B. napus* leaves (7dpt) were placed into the leaf chamber (9 cm^2^) at mid-day (12:00-14:00). Leaves were exposed to 200 µmol/m^2^/s and fed ambient air at a rate of 0.003 L/s [29]. Readings were recorded after 15 minutes and ten replicates, each replicate consisting of an individual *B. napus* seedling, were used to determine statistical significance between the mock and PA23-treated conditions with a Student’s t-test (P < 0.05)

## Results

### Phenotypic differences in *B. napus* growth

First, we examined the effects of PA23 on *B. napus* seedling development at 24 hpt and 7dpt. Plant size and dry weight of *B. napus* seedlings treated with PA23 were not significantly different compared to the mock-treated seedlings at 24 hpt (Figure 1A). However, at 7dpt the dry weight of PA23-treated seedlings increased by 56.8% (p = 0.0001) with a corresponding increase in the root/shoot ratio (p < 0.015) (Figure 1B).

**Figure 1.**
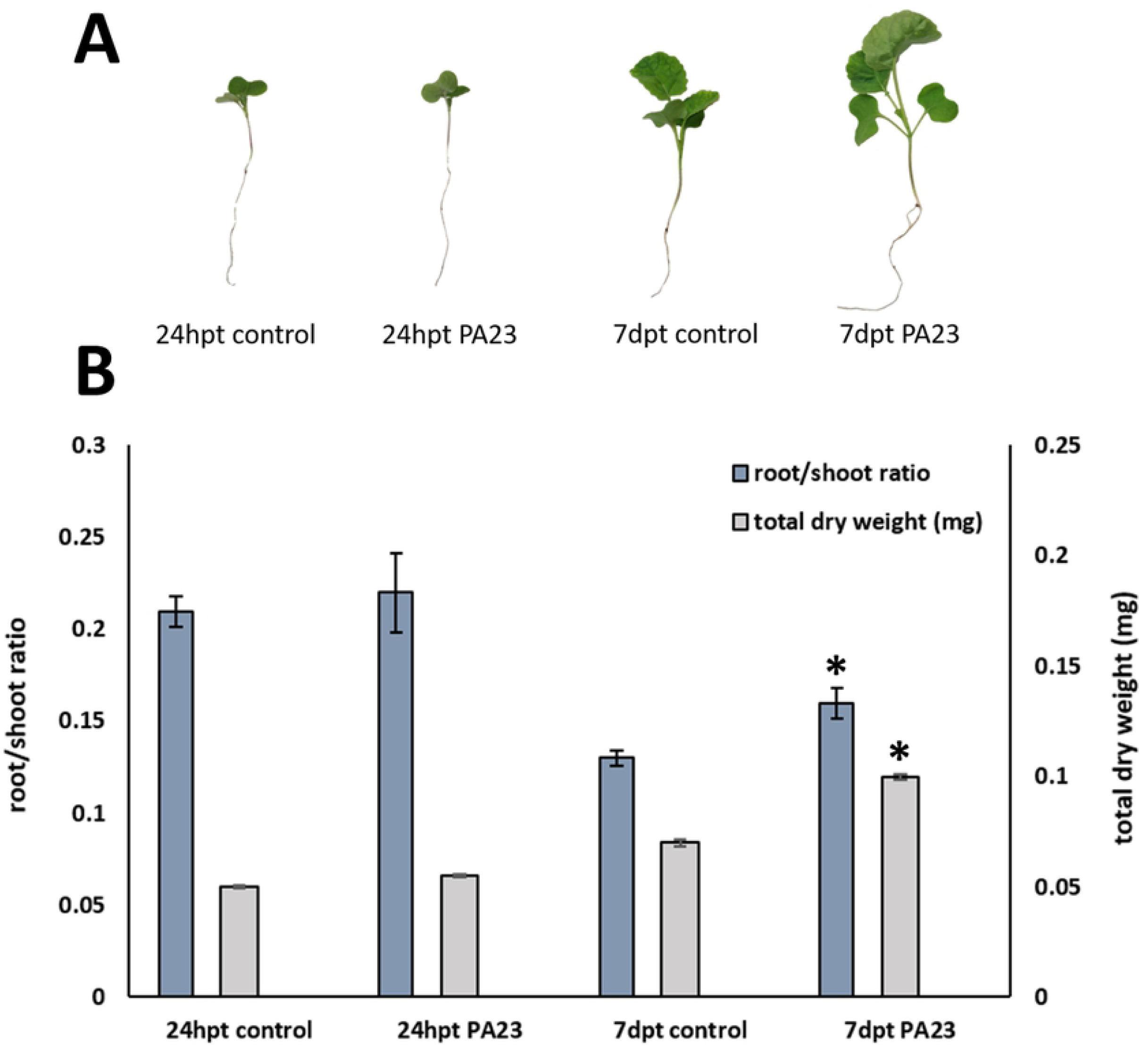
Development of *B. napus* seedlings at 24hpt and 7dpt. *B. napus* treated with lysogeny broth (mock-treatment) or a PA23 suspension (1×10^9^ cfu/mL. **A)** Representative *B. napus* seedlings showing phenotypic differences in the presence or absence of PA23. **B)** Total dry weight of *B. napus* and root/shoot ratios at 24hpt and 7dpt * indicates significance in a Student’s t-test (p < 0.05) and error bars represent standard error.

### Global comparisons of gene activity in the *B. napus* – *P. chlororaphis* PA23 interaction

To identify genes associated with increased growth in PA23-treated *B. napus* seedlings, we profiled the transcriptomes of the root and shoot tissue systems at 24 hpt and 7dpt. First, hierarchical clustering analyses revealed relationships between tissue systems and bacterial treatments. Samples first clustered by organ system followed by developmental time point, and finally by PA23 treatment (Figure 2A). Detected transcripts were then categorized into three expression levels: lowly (FPKM ≥1, <5), moderately (FPKM ≥5, <25), and highly (FPKM ≥ 25) expressed transcripts. We identified an even distribution of lowly (39%) and moderately (41%) expressed transcripts across all data points; however, we observed a 27% increase in highly expressed transcripts in PA23-treated shoots at 7dpt. Cumulatively, our dataset detected 56,508 transcripts with an FPKM value ≥ 1 or 56% of the *B. napus* genome. To identify genes contributing to increased plant growth, we conducted differential gene expression analyses of PA23-treated *B. napus* against mock-treated controls. At 24hpt, we detected 4,200 upregulated differentially expressed genes (uDEGs) in the shoot system and 3,267 uDEGs in the root system in response to PA23. The number of uDEGs decreased at 7dpt; 153 uDEGs were detected in the shoot system and 3,534 uDEGs were detected in the root system (Figure 3). Using these co-expressed gene sets, we performed GO term enrichment to identify putative biological processes associated with PA23 treatment.

**Figure 2.**
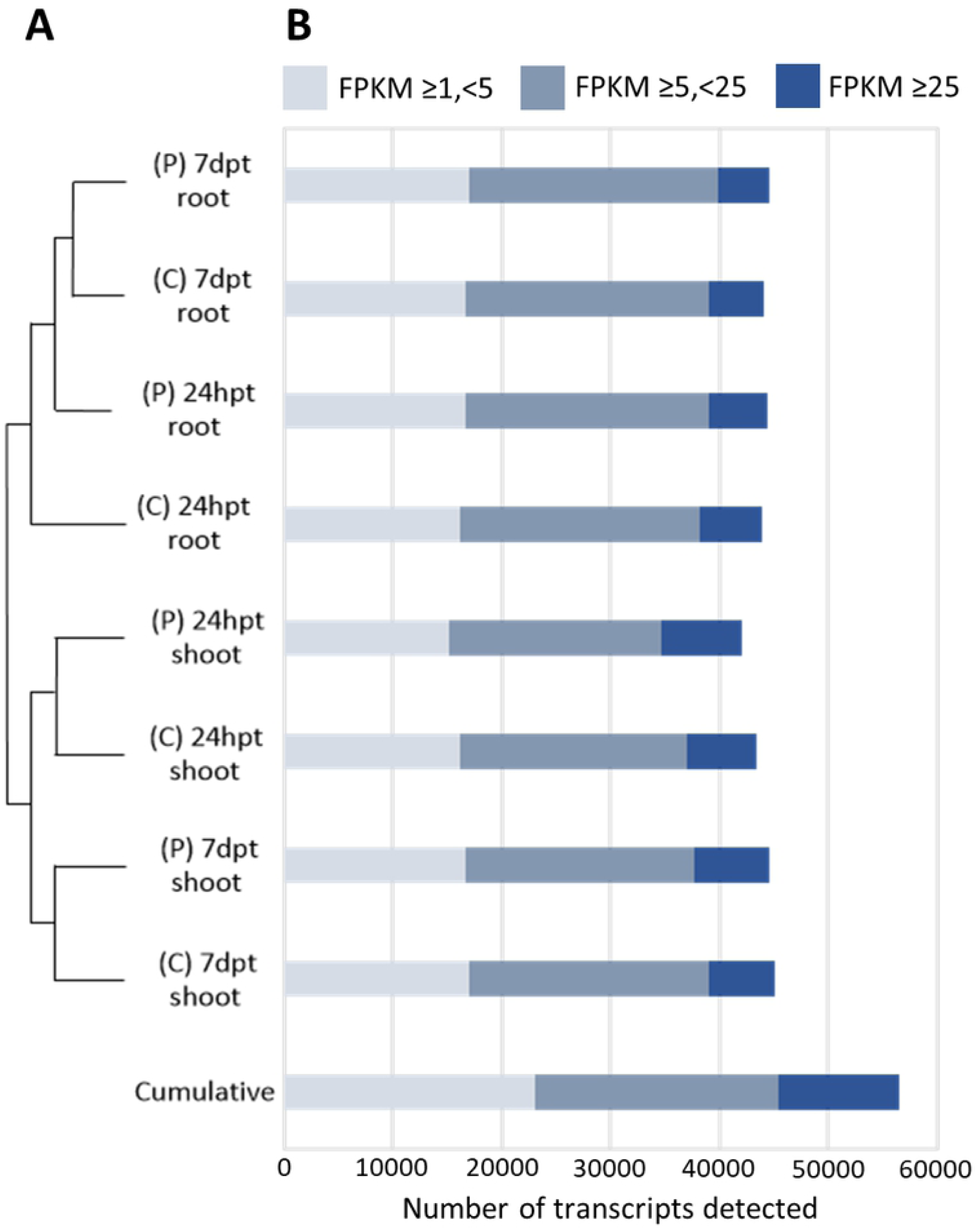
Hierarchical clustering and global gene activity in the *P. chlororaphis* PA23*-B. napus* interaction. **A)** Hierarchical clustering of all differentially expressed genes detected in the dataset. (P) indicates PA23 treatment and (C) indicates the mock-treatment group. **B)** Number of transcripts detected in both tissue systems across all treatments. Detected transcripts are subdivided into lowly (FPKM ≥1, <5), moderately (FPKM ≥5, <25), or highly (FPKM ≥ 25) expressed.

**Figure 3.**
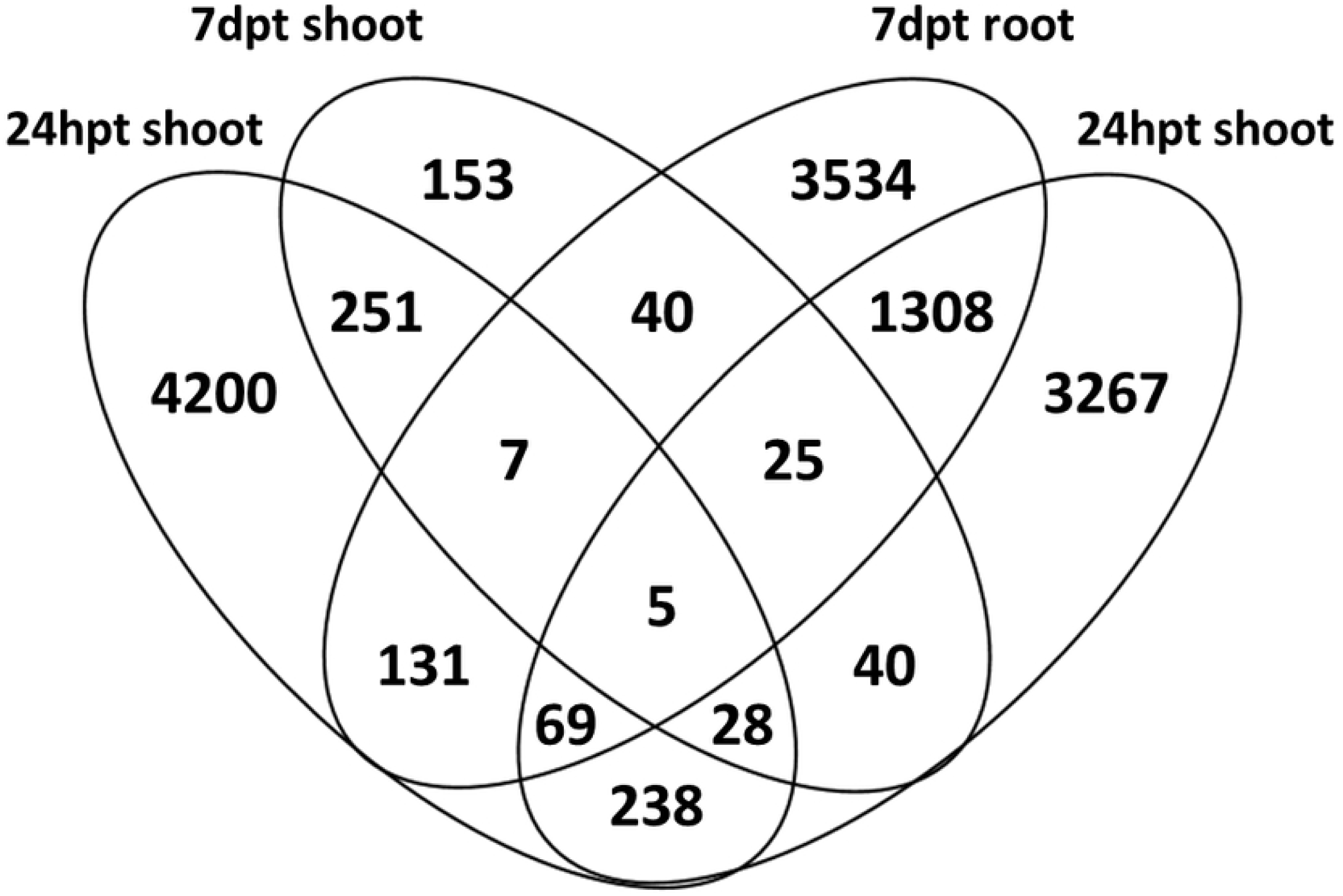
Differentially expressed genes identified in the PA23-treated *B. napus* seedlings in the root and shoot systems at 24hpt and 7dpt. Upregulated differentially expressed genes with overlapping regions showing the number of shared genes between tissue types and time points.

### PA23 increases expression levels of photosynthetic and reactive oxygen scavenging genes

Transcripts associated with the photosynthetic components were identified with the uDEGs in PA23-treated shoots at 24hpt and 7dpt. We also identified genes associated with reactive oxygen species (ROS) scavenging including superoxide dismutase activity and glutathione transferases. Specifically, we identified an upregulation of the photosystem gene, *LIGHT HARVEST COMPLEX PHOTOSYSTEM II SUBUNIT 6*, with an average increase of 28% and 39% in PA23-treated shoots at 24hpt and 7dpt, respectively. We investigated additional transcripts associated with photosynthesis and observed an upregulation in *PHOTOSYSTEM II SUBUNIT Q-2* (25%), *LIGHT HARVESTING CHLOROPHYLL PROTEIN COMPLEX II SUBUNIT B1* (29%), *ATP SYNTHASE SUBUNIT* (27%), *PHOTOSYSTEM I REACTION CENTER SUBUNIT PSI-N* (27%), AND *PHOTOSYSTEM I SUBUNITS* (average of 23%) against the mock treatment at 24hpt. Differential upregulation of photosynthetic genes shifted to pigment binding proteins at 7dpt; here we detected a 36% increase in transcript abundance associated with *CHLOROPHYLL A/B BINDING PROTEIN* in PA23-treated *B. napus* (Data in S2 Table).

Consistent with increases in the activity of photosynthetic genes, we observed an enrichment for catalase activity and superoxide dismutase activity, which are both associated with ROS scavenging in PA23-treated shoots at 24 hpt. To further characterize genes responsible for ROS quenching at 24 hpt, we investigated the levels of several transcripts including *COPPER/ZINC SUPEROXIDE DISMUTASE*, which increased by 26% in the shoots of PA23-treated *B. napus*. We observed an increase of *COPPER CHAPERONE* transcripts in PA23-treated shoots at both 24hpt (75% increase) and 7dpt (68% increase). To investigate the effects of PA23 on photosynthetic physiology, we extracted total pigment from PA23-treated and mock-treated tissue. We recorded an increase in chlorophyll a by 65% and chlorophyll b by 139% within 24 hours post treatment compared to untreated control tissues. At 7dpt, pigment levels accumulated to levels of 32% and 34% chlorophyll a and chlorophyll b, respectively in PA23-treated *B. napus* when compared against the untreated control (Figure 4A). Lastly, we measured the relative photosynthetic rates of PA23-treated and mock-treated *B. napus* seedlings at 7dpt to determine the relationship between transcript abundance, pigment concentration, and carbon fixation rates. Using IRGA and measuring the rate of CO_2_ depletion, we determined that PA23-treated *B. napus* plants exhibited a 60% increase in photosynthetic rates at 7dpt (Figure 4B). Taken together, *P. chlororaphis* PA23 increased the pigment concentration and photosynthetic rates of *B. napus* seedlings, which contribute to the observed growth phenotype.

**Figure 4.**
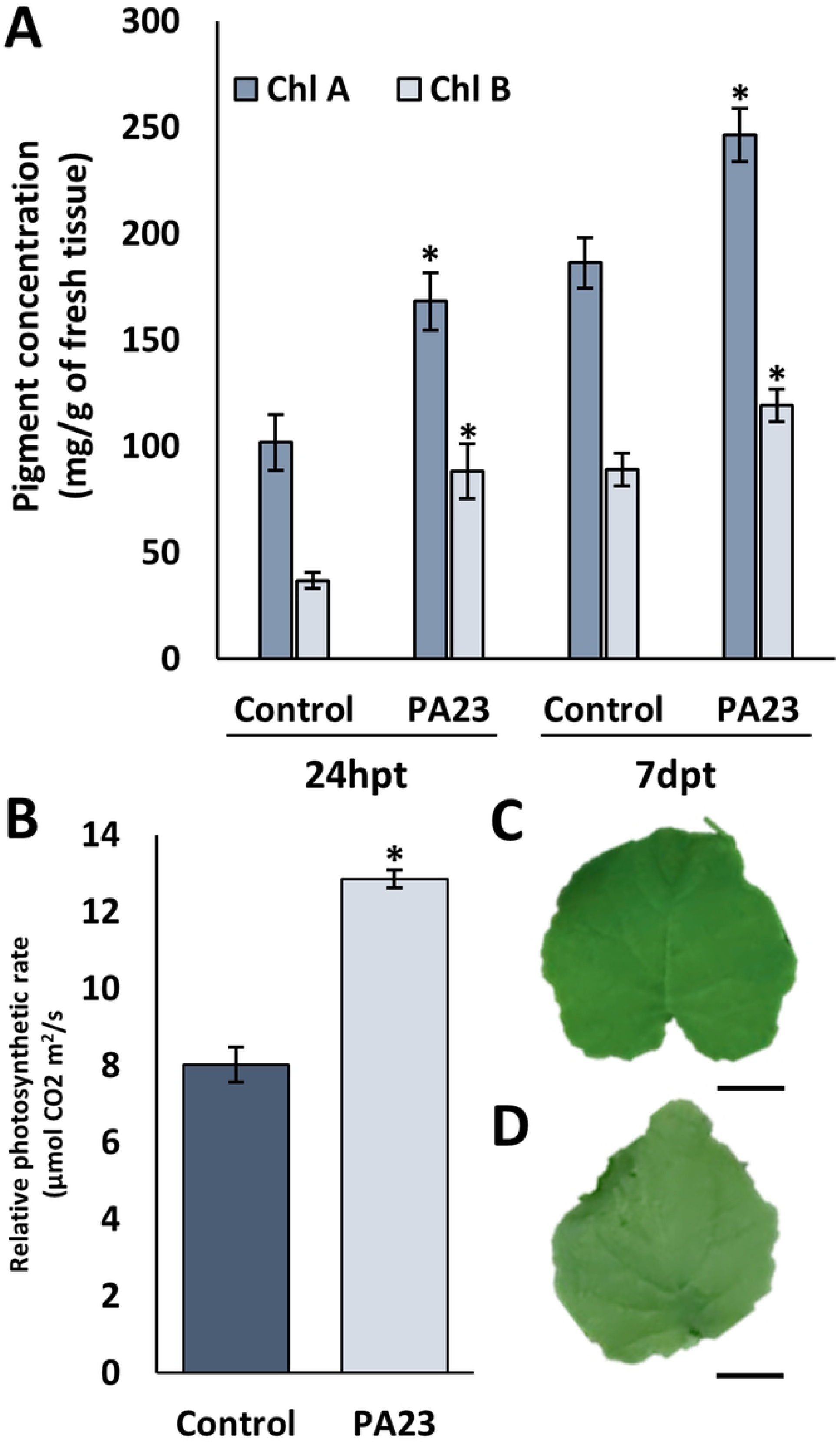
A)Pigment concentration of PA23-treated and mock-treated leaves at 24hpt and 7dpt. Absorbance values were measured at 663 nm and 645 nm and used to determine concentrations which were normalized to leaf input weight. * indicate statistical significance in a Student’s t-test (p < 0.05) and error bars represent standard error (n=15). **B)** Relative photosynthetic rates determined with IRGA equipment and CO_2_ depletion. * indicate statistical significance in a Student’s t-test (p<0.05) and error bars represent standard error (n=10). **C)** PA23-treated leaf at 7dpt, Scale bar = 1 cm. **D)** Control treated leaf at 7dpt, Scale bar = 1 cm

### PA23 promotes root hair development and nutrient transporter gene activity

At the root interface between the plant and bacteria, we identified an enrichment of GO terms associated with nutrient uptake (Figure 5A). For example, we observed an increase in the abundance of *AMT* (ammonium methyl transporter), *NRT* (nitrate transporter), and *PHT* (phosphorus transporter) transcripts in PA23-treated roots at 7dpt. Lastly, we discovered an enrichment of GO terms associated with cellular growth including cell wall biogenesis, cellulose synthase, and root hair elongation (Figure 5A). Specifically, we identified an upregulation of the essential root hair development gene – *ACTIN2*, which showed 12% and 21% increased transcript abundance at 24hpt and 7dpt, respectively (Data in S2 Table). To relate transcript abundance with phenotypic changes we used the model species -*Arabidopsis thaliana*, a close relative of *B. napus* in the Brassicaceae family. Microscopic visualization of PA23-treated and mock-treated roots at 7dpt revealed differences in root hair abundance. Specifically, the mock treatment (0.9% saline) exhibited a lower abundance of root hairs (Figure 5B). Whereas, PA23-treated plants had a higher abundance of root hairs along the primary root axis (Figure 5C).

**Figure 5.**
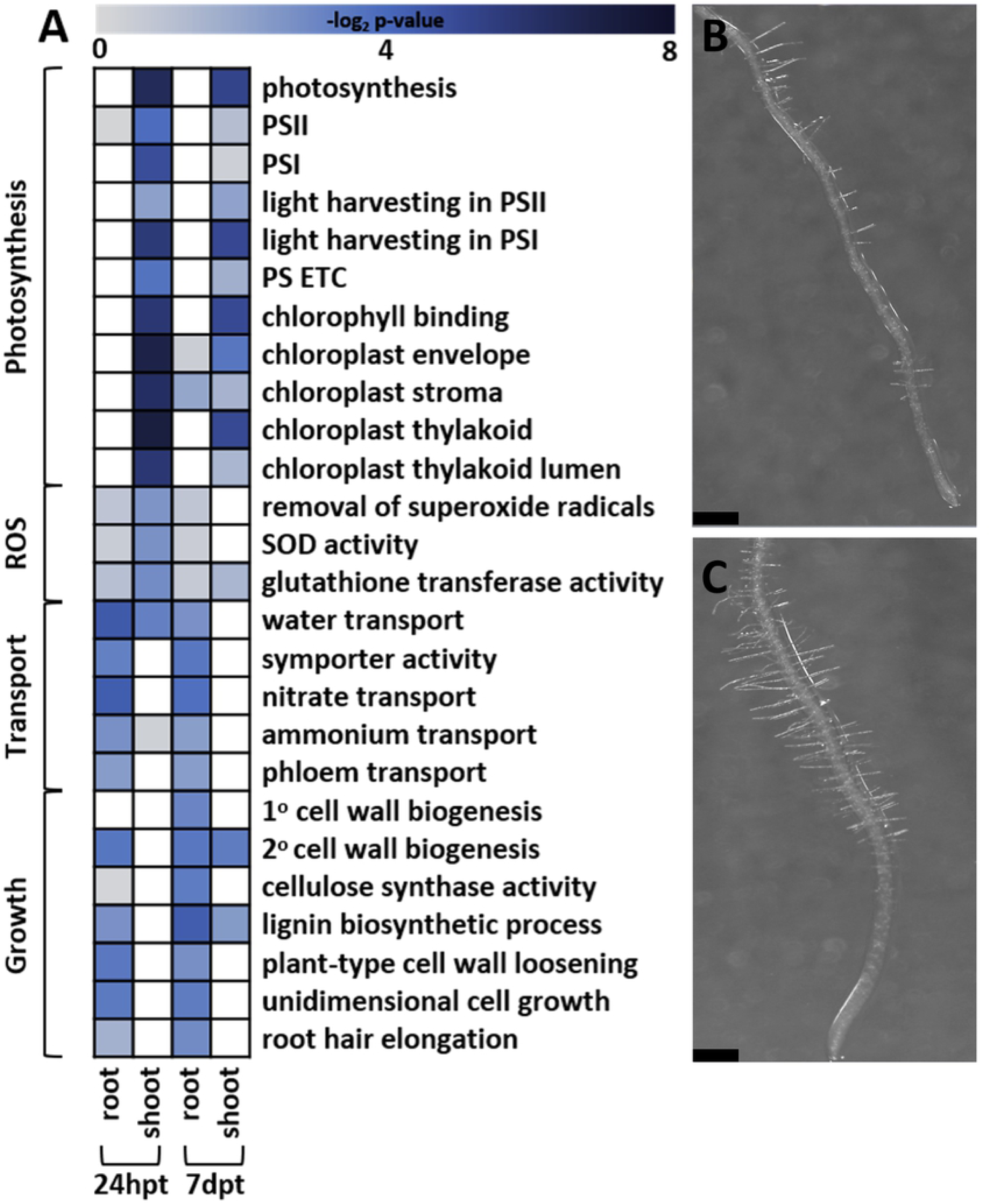
A)Heatmap of enriched Gene Ontology terms identified from upregulated genes with SeqEnrich. Darker colour represents a greater statistical enrichment **B)** *Arabidopsis thaliana* root imaged with a Leica M80 seven days after treatment with 0.9% saline. **C)** *Arabidopsis thaliana* root imaged with a Leica M80 seven days after treatment with 2000 cfu of *P. chlororaphis* PA23. Scale bar = 1000 µm. A complete list of enriched GO terms can be found in supplemental file 1.

### PA23 enhances plant growth through the modulation of phytohormone signalling

Next, we investigated genes associated with cytokinin, gibberellic acid, and auxin signalling. Roots exposed to PA23 were enriched for GA, CK, and IAA response genes at 24hpt and 7dpt. Notably, all transcripts related to GA response at 24hpt and 7dpt were homologs of the sucrose catabolism gene, *GLYCOSYL HYDROLASE FAMILY 32*, and their respective FPKM values increased by 43% and 86%. Genes associated with IAA signalling in the *AUXIN-INDUCED IN ROOT CULTURES* family were highly upregulated, with a 465% increase in PA23-treated roots at 24hpt (Data in S2 Table). Furthermore, multiple homologs of the CK/IAA response integrator, *SHY2*, were upregulated in the roots of PA23-treated *B. napus* with a 54% and 9% increase at 24hpt and 7dpt respectively. Additionally, we measured the production of IAA by *P. chlororaphis* PA23 using colorimetric spectrophotometry. Bacterial cultures were grown in the presence of L-tryptophan and we determined that *P. chlororaphis* PA23 produces IAA at a rate of 44.2 µg/hour (Figure B in S1 Text).

## Discussion

Crop production is of growing global concern and recent interest in the integration of environmentally friendly agricultural adjuncts has highlighted the potential use of PGPBs [30,31]. Currently, the mechanisms involved in growth promotion are not well understood.

Building on our understanding of the genetic mechanisms that underlie PGPB-induced plant growth is a critical step for the acceptance of PGPBs by regulatory agencies and the agricultural sector [1]. Our study aims to contribute to the growing body of information on the genetic mechanisms driving PGPB induced plant growth. Specifically, we investigated the mechanisms mediating enhanced growth during the *B. napus* – *P. chlororaphis* PA23 interaction. Our study revealed both molecular and phenotypic changes in the shoot and root systems of *B. napus* treated with *P. chlororaphis* PA23. In the shoots, we identified increased concentrations of the major photosynthetic pigments – chlorophyll a and chlorophyll b in the leaf tissue of *B. napus* treated with *P. chlororaphis* PA23. At the transcript level, our dataset identified increased abundance of photosynthetic transcripts such as *LHCB1*.*3*. This gene encodes a pigment binding protein of light harvesting complex II, which is responsible for light capture in the antenna complex of plants [32]. In an earlier study, mature canola plants treated with PA23 exhibited a 22% increase in total chlorophyll compared to untreated plants, which was accompanied by significant downregulation of the chlorophyll degradation pathway *CHLOROPHYLLASE 1* [15].

Other studies have identified changes in the expression of photosynthetic genes following PGPR treatment. For example, treatment of *Triticum aestivum* with *Dietzia natronolimnae* STR increased the concentrations of chlorophyll a, chlorophyll b, and carotenoids in photosynthetic tissues. Furthermore, PGPR treated *T. aestivum* were larger in size, which was attributed to the increased pigment concentrations and photosynthetic rates [33]. The expression of photosynthetic genes, such as *LHCB1*.*3*, is influenced by abiotic and biotic factors [34]. For example, plants acclimatize to changing light conditions through phytohormone signalling and differential expression of genes associated with photosynthesis. In *Hordeum vulgare*, the presence of exogenous cytokinin increased the expression rates of genes linked to chloroplast development and differentiation [35]. Tahir et al identified an increase in the expression rates of phytohormone signalling genes and enhanced photosynthetic rates in *Solanum lycopersicum* following treatment with *Bacillus subtilis* [36]. These changes were due to the volatile organic compounds produced by *B. subtilis* which altered gene expression in *S. lycopersicum*. We identified a similar increase in photosynthetic rates of *B. napus* following treatment with *P. chlororaphis* PA23. The production of cytokinin has been previously reported in *P. fluorescens* G20-18, which bolstered plant defense through the activation of cytokinin mediated immune responses [37]. Together, these studies suggest that bacterial metabolites can alter phytohormone signalling, photosynthetic transcript abundance, and photosynthetic rates. In the current study, we identified an enrichment of genes linked to cytokinin response in PA23-treated *B. napus*; however, we did not detect differential expression of genes associated with cytokinin biosynthesis. Thus, the expression patterns identified in our data and previous reports of volatile organic compound and cytokinin production in other *Pseudomonas spp*. suggest that *P. chlororaphis* PA23 may alter *B. napus* gene expression through a secondary or volatile compound [38,39]. Despite these findings, the identities of the entire suite of plant growth promoting metabolites produced by PA23 are unknown and further studies are required to elucidate the active metabolites produced by PA23.

Increased photosynthetic activity can have detrimental effects on cellular components and biological molecules. Under stress conditions such as drought, UV, and heat, ROS generation from the photosystems can increase [40]. The prevention of ROS mediated damage is facilitated by both non-enzymatic methods which absorb excessive light radiation, preventing the production of ROS, and enzymatic methods such as superoxide dismutases (SOD) that degrade cellular radicals [40]. These SODs are the first line of defense against ROS-mediated oxidation and SOD1 is one of the most potent cellular antioxidants [41]. In the current study, we identified enriched GO terms associated with ROS scavenging. For example, essential ROS scavenging genes such as *FSD 1, FSD III*, and *SOD* were upregulated in PA23-treated *B. napus* shoots 24hpt suggesting an increased demand for ROS scavenging. Under copper limited conditions, the expression of SODs are downregulated in *A. thaliana* and *Brassica juncea*, as copper is an essential component of the catalytic core of these enzymes [42]. Following PA23-treatement, we observed increased expression of genes associated with copper binding, copper chaperones, and copper uptake in the root dataset. Together, this may suggest that in response to increased SOD activity, *B. napus* requires elevated rates of copper sequestration to replenish depleted copper pools to counteract heightened levels of ROS production from the photosystems. *FSD I* is a cytoplasmic antioxidant that is preferentially expressed in *A. thaliana* under copper deficient conditions while *FSD III* is essential for chloroplast development.

Mutants deficient in *FSD III* are smaller in size, lack photosynthetic pigments, and are sensitive to oxidative stress [43]. Both *FSD I* and *FSD III* transcripts were more abundant in PA23-treated *B. napus*, further suggesting an increase in antioxidant demand. Together, the expression patterns of *SOD, FSD I*, and *FSD III* suggest that enhanced photosynthetic capabilities of PA23-treated *B. napus* is managed through increased expression rates of genes associated with antioxidants.

Growth regulation and patterning of the root is governed by concentration gradients of auxin and cytokinin. Crosstalk between these phytohormones is mediated by the signal integrator – SHY2, a transcriptional inhibitor of auxin response elements which regulate cell division [44]. The *ARR* (Arabidopsis response regulator) gene family is a group of cytokinin-induced transcription factors which increase expression rates of *SHY2* [45,46]. In contrast, auxin promotes the degradation of SHY2 and the activation of auxin response factors which increase transcriptional rates of auxin response elements resulting in elevated rates of cellular division. Thus, phytohormone concentration gradients and SHY2 work to balance cellular differentiation and division in the developing root. The production of auxin has been previously reported in other *Pseudomonas spp*. and PA23 carries the essential genes *TRYPTOPHAN 2-MONOOXYGENASE* and *INDOLE ACETMIDE HYDROLASE* for auxin biosynthesis via the indole acetamide pathway [21,47,48]. Further, we quantified the production of auxin by *P. chlororaphis* PA23 in liquid culture and observed moderate production of auxin or auxin homologs. Thus, the observed expression patterns of auxin response genes and *SHY2* in the root would suggest that growth promotion in the *B. napus* – *P. chlororaphis* PA23 interaction could be partially attributed to the presence of exogenous auxin. However, further studies characterizing bacterial products, such as auxin or other growth hormones, in the presence of developing plants are required.

The process of root hair development is separated into three phases: bulge formation, slow tip growth, and rapid tip growth [49]. Root hair extension occurs through asymmetric growth of the cytoskeletal element - actin. Application of chemicals that interfere with actin formation results in abnormal root hair initiation and early termination [50–52]. Specifically, the interruption of *DER1* in *A. thaliana* results in the absence of root hair on the primary and lateral roots [53]. The *der1*^-^ phenotype was rescued with an *ACT2* insertion under its native promoter [53]. Together, these studies suggest that *ACT2* is vital for root hair initiation and elongation. As previously described, nutrient availability and uptake are some of the most significant factors governing plant growth [54]. PGPRs can increase soil nutrient content and enhance nutrient uptake rates. For example, wheat treated with *P. fluorescens* under nutrient limited conditions produced similar yields when compared to optimally fertilized conditions, suggesting that *Pseudomonas spp*. can mobilize organic and inorganic nutrients [55]. Currently, the nutrient mobilization effects of *P. chlororaphis* PA23 and its effects on plant nutrient uptake is unknown. Nitrogen is an essential macronutrient and is primarily assimilated as nitrate. The uptake of nitrates occurs at the root hairs through two systems: high affinity transporters (HATs) and low affinity transporters (LATs) [56]. HAT expression is rapidly increased following the detection of increased soil nitrate levels; this response is accented in *A. thaliana* exposed to nitrate after long term nitrogen deficiency. Furthermore, *nrt2*.*1*^-^ assimilated significantly lower levels of nitrogen, suggesting that HATs are essential for rapid and efficient nitrogen uptake [57].

Ammonium is another source of nitrogen and uptake is facilitated by ammonium transporters (AMTs) at the root hairs. *A. thaliana amt1*.*1*^*-*^ *amt1*.*3*^*-*^ double mutants absorbed 70% less ammonium, while *AMT1*.*3* overexpression increased ammonium influx by 30% [58]. Together, these findings suggest that AMTs are essential for the uptake of ammonium and overexpression of *AMTs* enhance uptake rates. In the current study, we identified higher levels of *ACT2* transcript abundance in PA23-treated *B. napus* and higher occurrence of root hairs on the primary and lateral roots of PA23-treated *A. thaliana*. Further, we detected increased expression levels of *NRTs* and *AMTs* in our dataset. Together, these findings suggest that PA23 increases the abundance of nutrient transporters present on the root hairs which may increase nutrient uptake rates. Application to other field crops including *Glycine max*, the crop species from which PA23 was originally isolated, may result in similar increases in nutrient uptake which would decrease the dependence on chemical fertilizers.

In conclusion, our data suggests that *P. chlororaphis* PA23 is a potent PGPB that alters both the root and shoot systems of *B. napus*. Furthermore, our data identified that secretion of auxin by *P. chlororaphis* PA23 may be the primary contributor for plant growth promotion and potentially alter gene expression associated with photosynthesis in the shoot. However, our current understanding of the suite of metabolites produced by *P. chlororaphis* PA23 remains unknown and requires further investigation to identify potential active metabolites or small molecules. These metabolites could be the determining factor that governs the dual action of plant growth promotion and biocontrol activity exhibited by *P. chlororaphis* PA23. Several elements may contribute to this phenomenon including population density, quorum sensing, and differing host plant species. Further studies using quorum sensing and signalling mutants of *P. chlororaphis* PA23 could shed light on the underlying mechanism that determines its activity as a PGPB. Together, our study provides preliminary evidence for the mechanisms that drive enhanced plant growth by *P. chlororaphis* PA23, but further studies are required to determine the mechanisms that control metabolite production and biocontrol or plant growth promotion activity.

## Acknowledgements

Funding was provided by the Natural Science and Engineering Research Council to MFB, TRdK, and WGDF through a Collaborative Research and Development Grant and an NSERC Vanier Scholarship to MGB. The authors thank Kirsten Biggar and Rachel Robinson for technical assistance.

